# Uncovering non-random binary patterns within sequences of intrinsically disordered proteins

**DOI:** 10.1101/2021.08.19.456831

**Authors:** Megan C. Cohan, Min Kyung Shinn, Jared M. Lalmansingh, Rohit V. Pappu

**Affiliations:** Department of Biomedical Engineering and Center for Science & Engineering of Living Systems (CSELS), Washington University in St. Louis, MO 63130, USA; Department of Physics, Washington University in St. Louis, MO 63130, USA

**Author notes:** Equal contributions (Megan C. Cohan, Min Kyung Shinn).

## Abstract

Sequence-ensemble relationships of intrinsically disordered proteins (IDPs) are governed by binary patterns such as the linear clustering or mixing of specific residues or residue types with respect to one another. To enable the discovery of potentially important, shared patterns across sequence families, we describe a computational method referred to as NARDINI for **N**on-random **A**rrangement of **R**esidues in **D**isordered **R**egions **I**nferred using **N**umerical **I**ntermixing. This work was partially motivated by the observation that parameters that are currently in use for describing different binary patterns are not interoperable across IDPs of different amino acid compositions and lengths. In NARDINI, we generate an ensemble of scrambled sequences to set up a composition-specific null model for the patterning parameters of interest. We then compute a series of pattern-specific z-scores to quantify how each pattern deviates from a null model for the IDP of interest. The z-scores help in identifying putative non-random linear sequence patterns within an IDP. We demonstrate the use of NARDINI derived z-scores by identifying sequence patterns in three well-studied IDP systems. We also demonstrate how NARDINI can be deployed to study archetypal IDPs across homologs and orthologs. Overall, NARDINI is likely to aid in designing novel IDPs with a view toward engineering new sequence-function relationships or uncovering cryptic ones. We further propose that the z-scores introduced here are likely to be useful for theoretical and computational descriptions of sequence-ensemble relationships across IDPs of different compositions and lengths.

## Introduction

Approximately 30-40% of eukaryotic proteomes contain proteins that are either entirely disordered or include disordered regions ^1,2^. Under typical approximations of physiological solution conditions (pH 7.4, 37°C, and 100-300 mOsm of salts and solutes), intrinsically disordered proteins / regions (IDPs / IDRs) adopt heterogeneous ensembles of conformations, although some of these systems can adopt stable folds in 1:1 or higher-order complexes with their binding partners ^3^. Many IDPs / IDRs are characterized by significant sequence variations within orthologs and homologs ^4-6^. Although sequence identity and similarity tend to be poorly conserved across orthologous IDRs, one can compare amino acid compositions across orthologs ^7-10^. This often reveals the conservation of compositional biases that might point to conserved composition-function relationships ^9,10^. The relevant parameters include overall amino acid compositions ^7^, sequence lengths ^11^, net charge per residue ^12-14^, and the identification of functionally relevant short linear motifs (SLiMs) ^15-20^. One example of a family of IDRs with a well-studied composition-to-function relationship is the disordered RGG domain found in proteins that drive phase separation ^21-23^. While the actual sequences cannot be aligned without inserting and extending numerous gaps, the overall compositional profiles are similar across this family of sequences ^22^.

Binary sequence patterns have also been established as important determinants of sequence-ensemble-function relationships of IDPs / IDRs ^4,5,8,9,24-28^. These are quantified using parameters for the segregation or mixing of a specific residue / residue type with respect to another type of residue ^29^ or all other residues in the sequence ^30^. Previous studies have demonstrated the functional importance of binary patterning parameters through mutational studies where sequence compositions were fixed, and the impacts of variations in linear sequence patterns on functions were assessed ^20,24,26,30-32^. Studies highlighting the importance of binary patterns, have prompted comparisons of context-dependent features within sets of IDPs / IDRs as a route to inferring potentially important sequence features. For example, Buske et al., noted that despite having different compositions, the extent of segregation vs. mixing of oppositely charged residues within IDR sequences, as quantified in terms of the parameter (κ), was bounded between 0.15 and 0.4 for the intrinsically disordered C-terminal linkers (CTLs) of FtsZs ^33^. Here, we show that the original binary patterning parameters, and generalizations of such parameters are not interoperable for quantitative comparisons across different compositions. Specifically, if amino acid compositions are different from one another, then a specific κ value does not imply similar degrees of segregation or mixing of oppositely charged residues. To remedy this deficiency and enable the extraction of significant binary patterns within IDRs, we developed and deployed NARDINI (**N**on-random **A**rrangement of **R**esidues in **D**isordered **R**egions **I**nferred using **N**umerical **I**ntermixing), an algorithm that is available as an extension of localCIDER ^28^, a freely available codebase for analyzing physico-chemical properties of IDRs. We show, using a set of examples, that the patterns we extract are consistent with known sequence-function relationships in specific systems.

## Results

### Parameters that quantify extents of segregation vs. mixing of different types of residues vary with amino acid composition and sequence length

A parameter that has been used to quantify sequence-ensemble-function relationships of IDRs is κ (which we will denote here as κ_+–_) ^29^. This parameter, which was introduced to quantify the overall segregation vs. mixing of oppositely charged residues with respect to one another ^9,24,29^, is defined as follows: for a given IDR, we tally *f*_+_ and *f*_–_, the overall fractions of positive and negatively charged residues, respectively. This is then used to compute the overall charge asymmetry (σ_+-_) using: 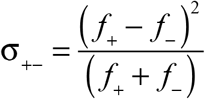. A sliding window comprising *g* residues is defined, and within each window *j*, we compute the local charge symmetry 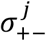. The mean squared deviation between the local and global charge asymmetry is computed as: 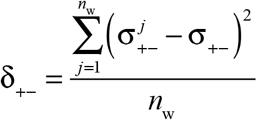, where *n*_w_ denotes the number of sliding windows used in the calculation. The mean squared deviation is normalized using a statistical model to estimate the maximal possible deviation 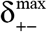 that is realizable for the composition such that 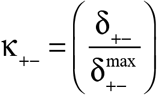, thereby ensuring that 0 ≤ κ_+–_ ≤ 1. The original normalization was designed to ensure that a specific value of κ_+–_ measures similar extents of segregation vs. mixing of oppositely charged residues across IDRs of different compositions. The validity of the normalization and interoperability of normalized values was tested using a set of sequences with high fractions of charged residues. We asked if values of κ_+–_ vary or stay fixed across the full range of FCR values for a fixed composition. Here, FCR is the fraction of charged residues defined as (*f*_+_ + *f*_–_) ^28^.

We considered sequences comprising three residues A, E, and K. We fixed the net charge per residue (NCPR) of the sequences to be zero, i.e., (*f*_+_ – *f*_–_) = 0. For a given sequence length, set here to be 50 or 100, and fixed composition achieved by fixing the values of *f*_+_, *f*_–_, and *f*_A_ such that (*f*_+_ + *f*_–_ + *f*_A_) = 1, we generated 10^5^ randomly shuffled sequence variants. For each sequence variant, we calculated the value of κ_+–_. For FCR values greater than 0.1, the distribution of κ_+–_ values could be fit using a gamma distribution (**Figure S1**). The mean (*m*) of the gamma distribution is the most likely value of κ_+–_ for the specific composition. We asked if the value of *m* depends either on FCR or sequence length. The impact of FCR was assessed by repeating the calculations described above for different values of FCR, operating under the constraints of NCPR = 0 and (*f*_+_ + *f*_–_ + *f*_A_) = 1.

Figure 1 shows how *m*(κ_+–_) varies with FCR for sequences that are 50 and 100 residues long. The length dependence is minimal and the relative flatness of *m*(κ_+–_) as a function of FCR for higher fractions of charged residues (FCR > 0.6) suggests that the original normalization is reasonable for certain categories of sequences. However, the most likely value for κ_+–_ does vary with amino acid composition when FCR is below 0.6. This is sub-optimal since a majority of IDPs tend to have FCR values that are ≤ 0.3 ^9^. Hence, while κ_+–_ is suitable for comparing the extent of segregation vs. mixing of oppositely charged residues across sequence variants that have identical compositions and lengths ^9,24-26,29^ the original normalization ^29^, which works well for sequences with FCR above 0.6, is inadequate for enabling comparisons across sequences with different compositions, especially when the FCR is below than 0.6.

**Figure 1.**
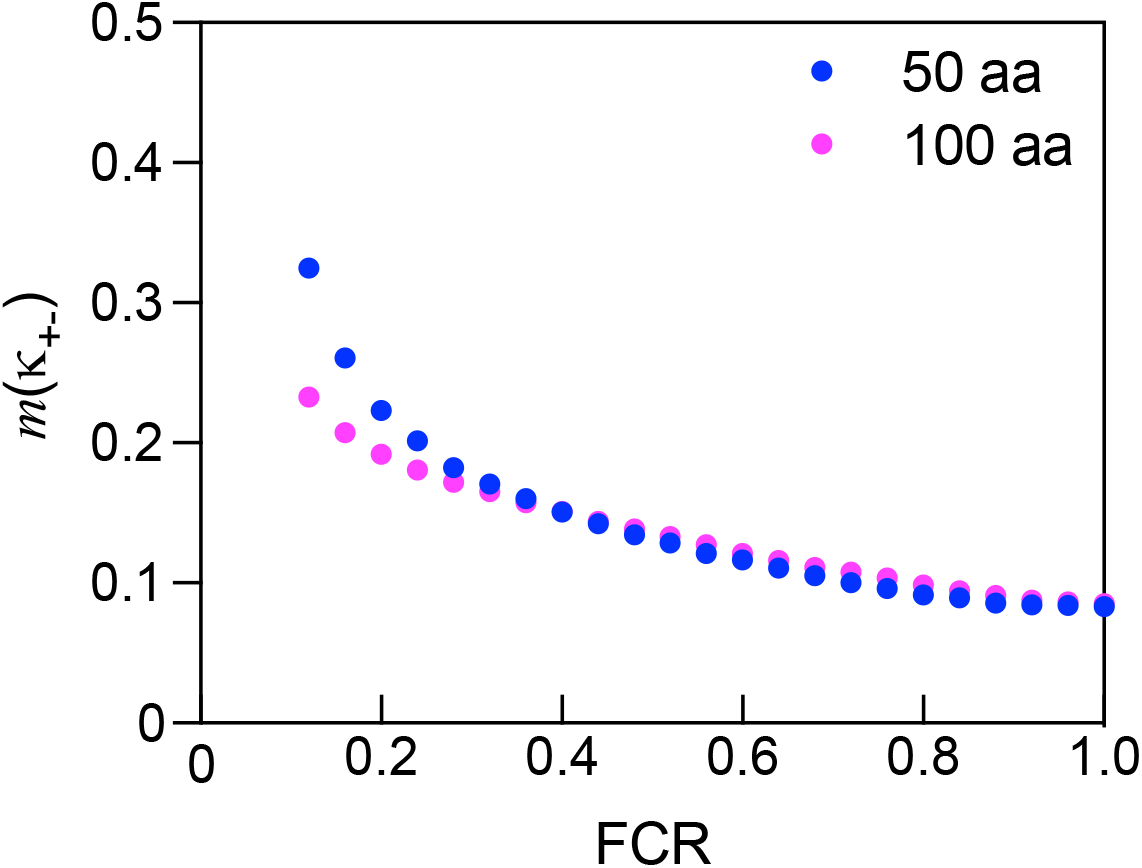
Plot of how the most likely value of κ_+–,_ measured as the mean of the gamma distributions (*m*(κ_+-_), depends on the fraction of charged residues (FCR) for sequences of 50 and 100 residues. The null-scramble expectations, i.e., the mean values of gamma distributed κ_**+–**_ values are shown for sequences that 50 and 100 residues long as a function of FCR. The mean value of κ_**+–**_ is dependent on FCR for low values of FCRs (< 0.3) and this dependency is also manifest for different sequence lengths.

Alternative parameters have been proposed and deployed to quantify the linear mixing vs. segregation of oppositely charged residues. These include the sequence charge decoration (SCD) parameter of Sawle and Ghosh ^34^. The dependence on composition that we observe for SCD (**Figure 2**), is even more pronounced for SCD than for κ_**+–**_. Further, is also a clear length dependence to the most likely value of SCD, which is not observed for κ_**+–**_.

**Figure 2.**
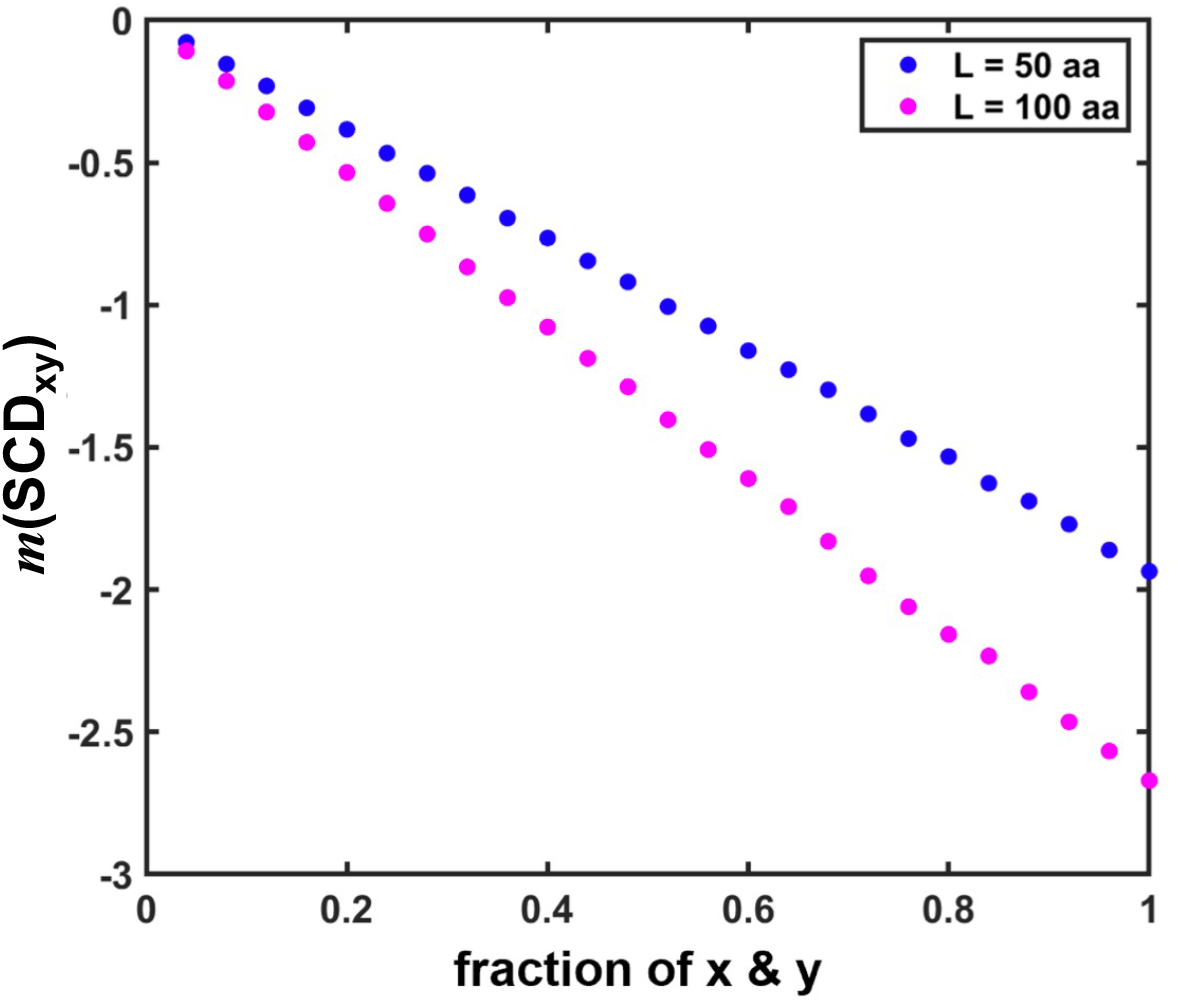
The mean value of SCD, quantified from the gamma distribution of SCD values for 10^5^ scrambled sequences, depends on the sequence composition and length. The null-scramble expectations of SCD are plotted for sequences of 50 and 100 residues as a function of fraction of residue x that is positively charged and y that is negatively charged (SCD_xy_). The expectation of SCD is dependent on the sequence length.

In addition to κ_**+–**_ and SCD, Martin et al.,^20,30^ introduced the Ω_x_-family of patterning parameters to quantify the linear mixing versus segregation of a residue x or residues of type x with respect to all other residues. Here, we find that the most likely values of Ω_x_ show a non-monotonic dependence on amino acid composition (**Figure 3**). Overall, we conclude, based on the analyses summarized in **Figures 1-3**, that despite their importance as descriptors of sequence-ensemble-function relationships ^9,20,24-26,28-30,35^, extant parameters that quantify mixing vs. segregation of different types of amino acids within IDRs cannot be deployed to compare patterns across sequences of different amino acid compositions and lengths. This creates a problem for uncovering patterns of potential functional importance, especially across hypervariable IDRs within orthologous systems. We solve this problem by deploying a new method, NARDINI, as described next.

**Figure 3.**
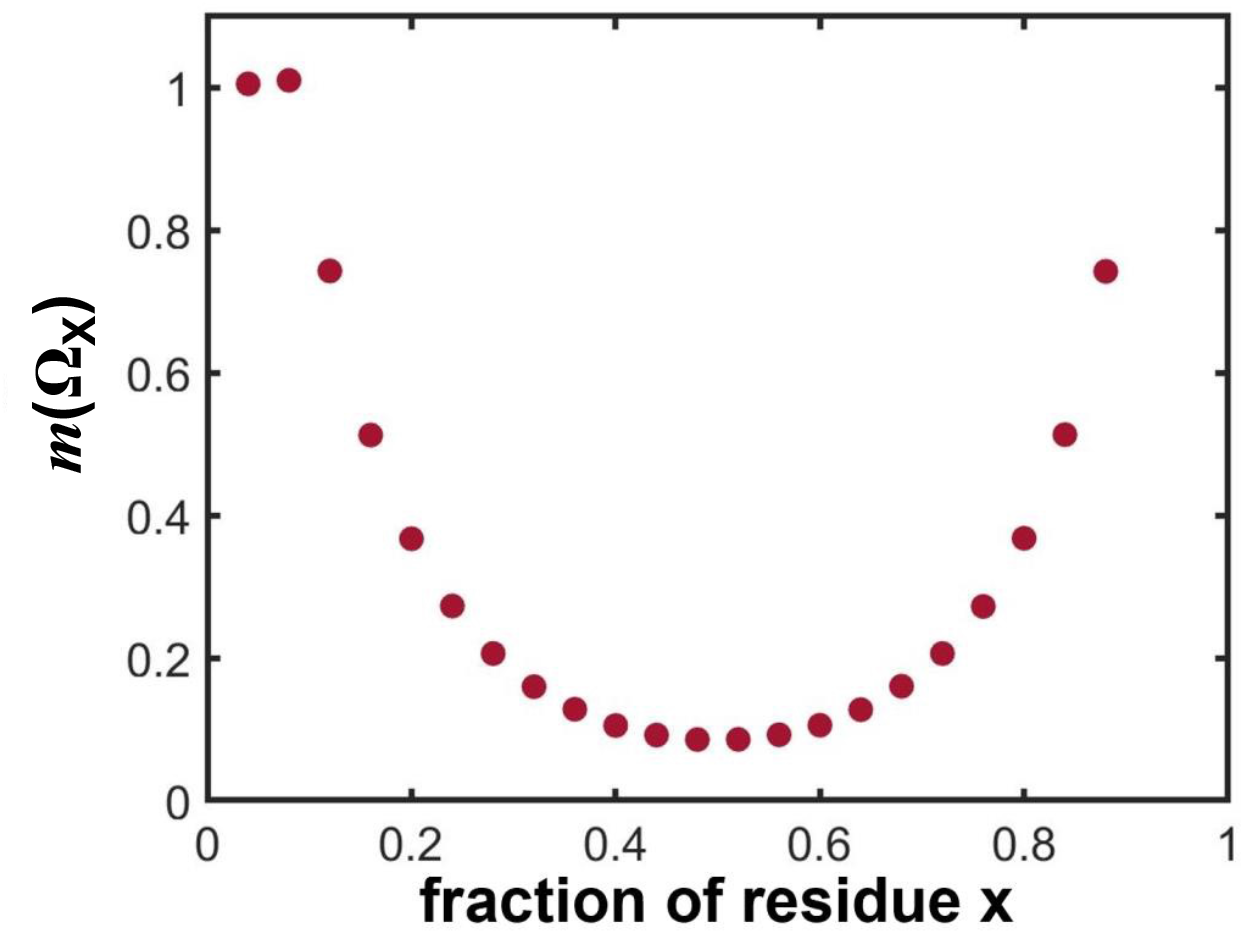
The mean value for the Ω parameter, as extracted from the gamma distribution obtained using 10^5^ randomly shuffled sequences, depends on amino acid composition. The plot shows the null-scramble expectations of the most likely value for (Ω_x_) for sequences of 100 residues as a function of the fraction of residue x.

### Generation of a null model

The first step of the NARDINI method is the generation of a null model. The workflow is summarized in **Figure 4**. For an IDR of interest, we generate 10^5^ independent scrambled sequences. In this approach, the amino acid composition is fixed, and the linear sequence is randomly shuffled. The patterning parameter of interest is calculated for each of the 10^5^ randomly generated sequences. For all parameters of interest, the random generation of sequence scrambles leads to maximum entropy-based gamma distributions. The mean (*m*) of the gamma distribution is the null expectation for the patterning parameter of interest. Next, for each sequence *i* in the set of scrambles, and patterning parameter of interest designated as *q*, we compute the sequence-specific z-score as: 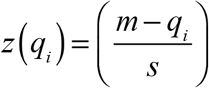. Here, *z*(*q*_*i*_) is the z-score of sequence *i* for the patterning parameter *q*, whereas *m* and *s*, respectively, are the mean and standard deviation of the distribution *P*(*q*). This is a gamma distribution of the patterning parameter *q* for sequences that have compositions and lengths that are identical to those of sequence *i*. For every parameter *q*, we can use the z-scores to identify patterns deemed to be statistically significant with respect to the null model. The z-scores enable two types of comparisons: We can quantify the extent to which the value of a specific patterning parameter deviates from random expectations, defined as z ≈ 0, since in this case, the parameter of interest does not deviate from the null expectation. Additionally, we can compare z-scores for the patterning parameter of interest across a set of IDRs from orthologous systems and gain a sense of the extent to which the pattern in question is statistically significant. Comparisons across different sequences are feasible because z-scores have a well-defined statistical interpretation, and for each sequence it is calibrated by composition and length-specific distribution of values for the patterning parameter of interest.

**Figure 4.**
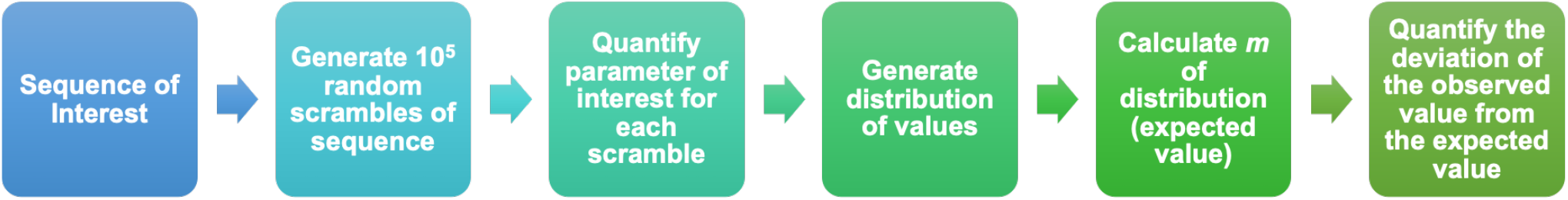
Workflow for the calculation of z-score matrices for various binary patterns. Process includes generating the null-scramble model (“null model”) and calculating the deviation of the observed value from the null model as z-scores.

### Prescribing a set of patterning parameters

There are two broad categories of parameters that quantify binary patterns within IDRs. We refer to these as the δ- and Ω-family of parameters. These are defined as follows: Consider a sequence defined by a specific amino acid composition such that *f*_x_ and *f*_y_ quantify the fractions of amino acids x and y, respectively. Further, (*f*_x_ + *f*_y_) ≤ 1. We define 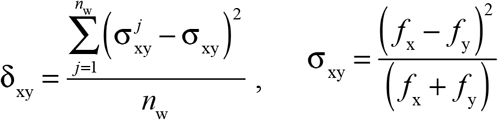, and 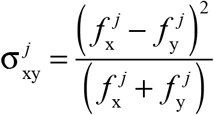 is the local asymmetry within window *j*. Here, we have replaced + and – from the previous section with x and y. Likewise, for *f*_x_ + *f*_y_ = 1, i.e., for a binary classification where residues either do or do not belong to group x, we can define the Ω-family of parameters using: 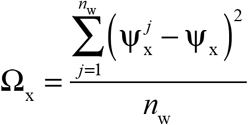, where 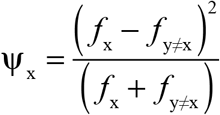 and 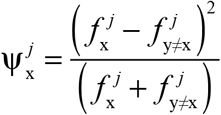. In both cases, *n*_w_ is the number of sliding windows across which the deviation between local and global asymmetries are calculated. For a sliding window of *g-*residues, *n*_w_ = *N – g+*1, where *N* is the sequence length. Following previous calibrations ^20,29,30^, we set *g* = 5.

The δ- and Ω-family of parameters are computed for specific groups of residues / residue types. We define eight such groups and denote these as polar residues μ ≡ {S,T,N,Q,C,H}, hydrophobic residues h ≡ {I,L,M,V}, basic residues + ≡ {R,K}, acidic residues – ≡ {E,D}, aromatic residues π ≡ {F,W,Y}, alanine A≡{A}, proline P ≡ {P}, and glycine G ≡ {G}. These groupings are based on similarities of stereochemistry, charge, aromaticity, size, and hydrophobicity. The groupings lead to the calculation of 36 distinct patterning parameters. These include eight parameters from the Ω-family *viz*., Ω_μ_, Ω_h,_ Ω_+_, Ω_–_, Ω_π_, Ω_A_, Ω_P_, and Ω_G_ and 28 parameters from the δ-family, where δ_xy_ represents every unique pairwise combination of {μ,h,+,– ,π,A,P, G}. For every IDR of interest, the NARDINI algorithm generates a z-score matrix, a schematic of which is depicted in **Figure 5**.

**Figure 5.**
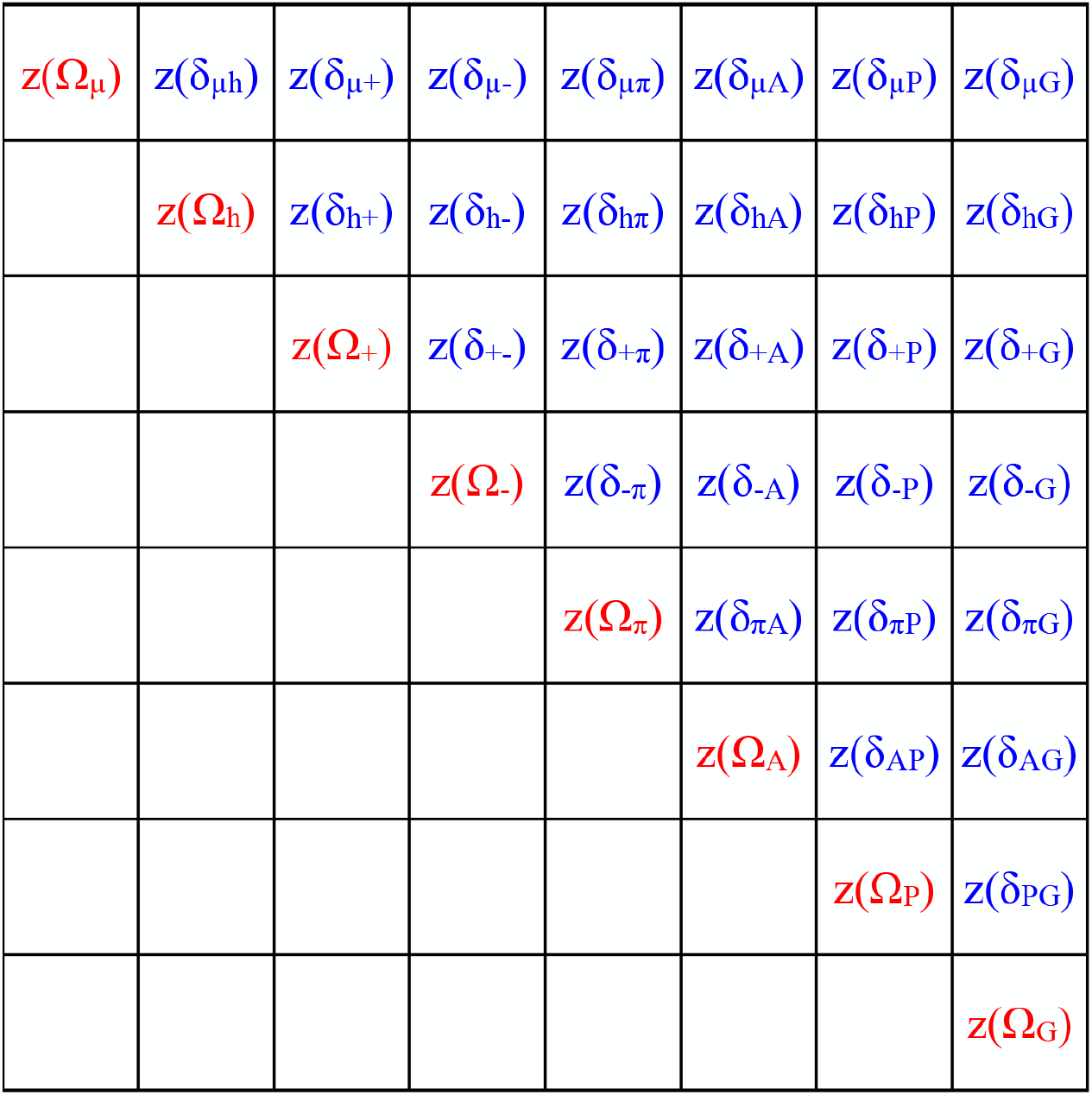
Elements of a typical, sequence-specific z-score matrix. Each cell quantifies the z-score of a specific patterning parameter. The diagonal elements are z-scores for Ω parameters whereas the off-diagonal elements are z-scores for the δ-parameters. Residues are grouped into eight categories: (μ) {S,T,N,Q,C,H}, hydrophobic; (h) {I,L,M,V}; positive (+) {R,K}; negative (-) {E,D}; aromatic (π) {F,W,Y}; alanine {A}; proline {P}; and glycine {G}.

We use an 8×8 matrix where only the diagonal and upper triangular values are considered. Note that the matrix size changes as we change the number of sets of residues / residue types of interest. The diagonal elements in the matrix are z-scores for different Ω-values that quantify the linear mixing versus segregation of a residue or a residue type with respect to all other residues. The off-diagonal elements in the matrix are z-scores for δ-values that quantify the linear mixing versus segregation of pairs of residues / residue types. When sequences contain fewer than 10% of a residue or residue type, all z-scores that involve that residue type are set to zero. Note that the z-score matrix is specific to a given sequence and helps identify potentially important binary patterns that are present in and IDP / IDR. We used sequence-specific z-score matrices to identify the Ω and δ parameters that have z-scores are greater than zero. Next, we illustrate the NARDINI approach by analyzing three distinct sequence families.

### Analysis of the prion-like low complexity domain from hnRNPA1

Prion-like low complexity domains (PLCDs) feature prominently among IDRs that are known to be drivers of phase separation and the formation of distinct types of biomolecular condensates ^19,36^. The PLCD from the RNA binding protein hnRNPA1 *homo sapiens* (referred to as the A1-LCD) has been used as an archetypal system to uncover sequence-specific driving forces for phase separation of PLCDs ^37^. A recent study showed that aromatic residues (Tyr / Phe) are the primary cohesive motifs (stickers) in the A1-LCD system ^20^. The linear segregation vs. mixing of aromatic residues contributes directly to the driving forces for phase separation and determines whether the condensates are liquid-like or amorphous solids ^20^.

We used the NARDINI approach to analyze the A1-LCD sequence (**Figure S2**). The z-score matrix shown in **Figure 6**, indicates the presence of four features that deviate significantly from the null model. First, the linear clustering of polar residues μ ≡ {S,T,N,Q,C,H} has a z-score for Ω_μ_ of +1.96. This derives from the presence of blocks of Ser residues along the sequence. Second, deviation from the null expectation for linear clustering is observed for Gly residues with the z-score of +3.03 for Ω_G_. Third, the Ser and Gly blocks tend to be segregated from one another, as evidenced by the z-score value of +3.93 for δ_μG_. And fourth, the z-score value for Ω_π_ is –1.86. This is consistent with the documented preference for uniform dispersion of aromatic residues along the linear sequence ^20^. The four patterns deemed to be statistically significant are consistent with features that have been shown to be directly relevant to the phase behavior of A1-LCD ^20,38-40^.

**Figure 6.**
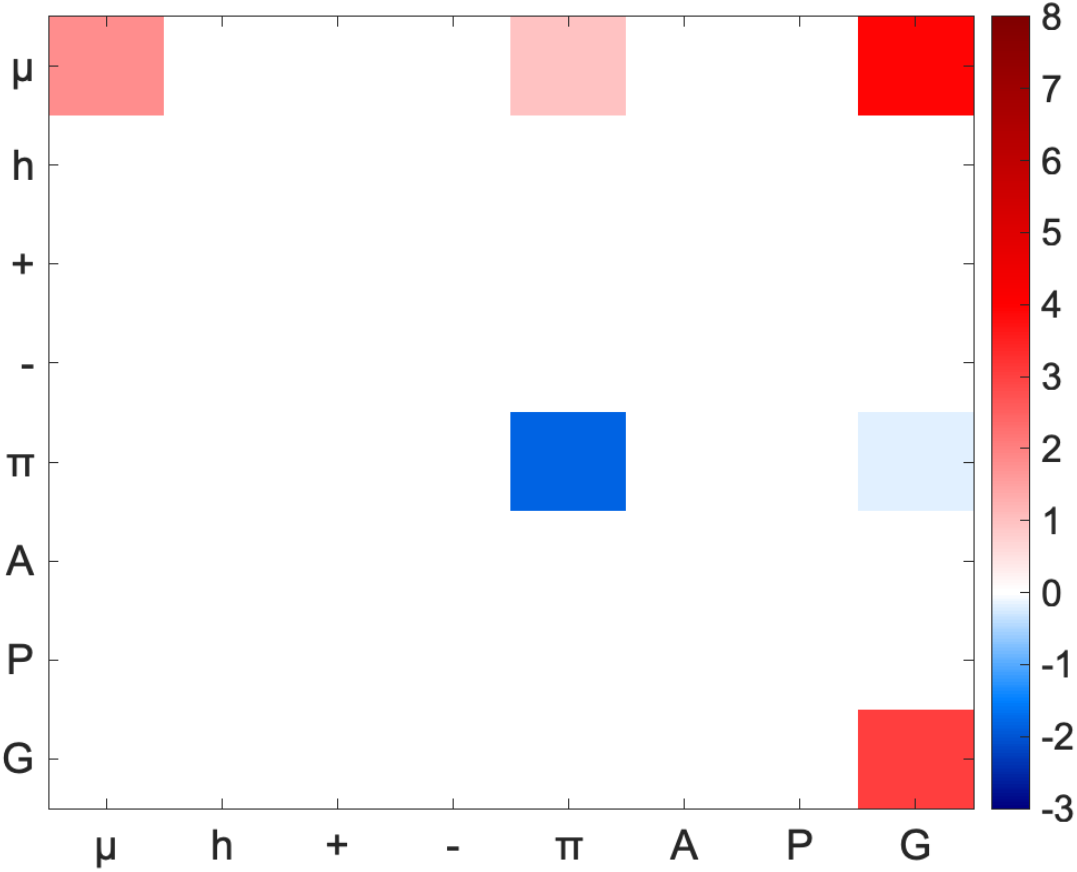
The z-score matrix of Al-LCD. A1-LCD shows non-random segregation of polar and glycine residues from one another and from other residues. This sequence also features non-random dispersion of aromatic residues. White squares on the checkerboard plot imply that the associated z-scores are ≈ 0.

Next, we analyzed z-score matrices across PLCDs derived from 849 homologs of hnRNP-A1 (**Figure 7**). For this analysis, we used the set of sequences was curated by Bremer et al.,^10^. Again, the salient binary patterns across the PLCDs are (i) the uniform dispersion of aromatic residues, (ii) the segregation of Gly and polar residues into distinct clusters, and (iii) the segregation of Gly and polar clusters from one another. These findings suggest that sequence features that contribute to phase separation of the A1-LCD system are qualitatively conserved, although the differences in z-scores suggest a quantitative titration of sequence features across homologs ^10^.

**Figure 7:**
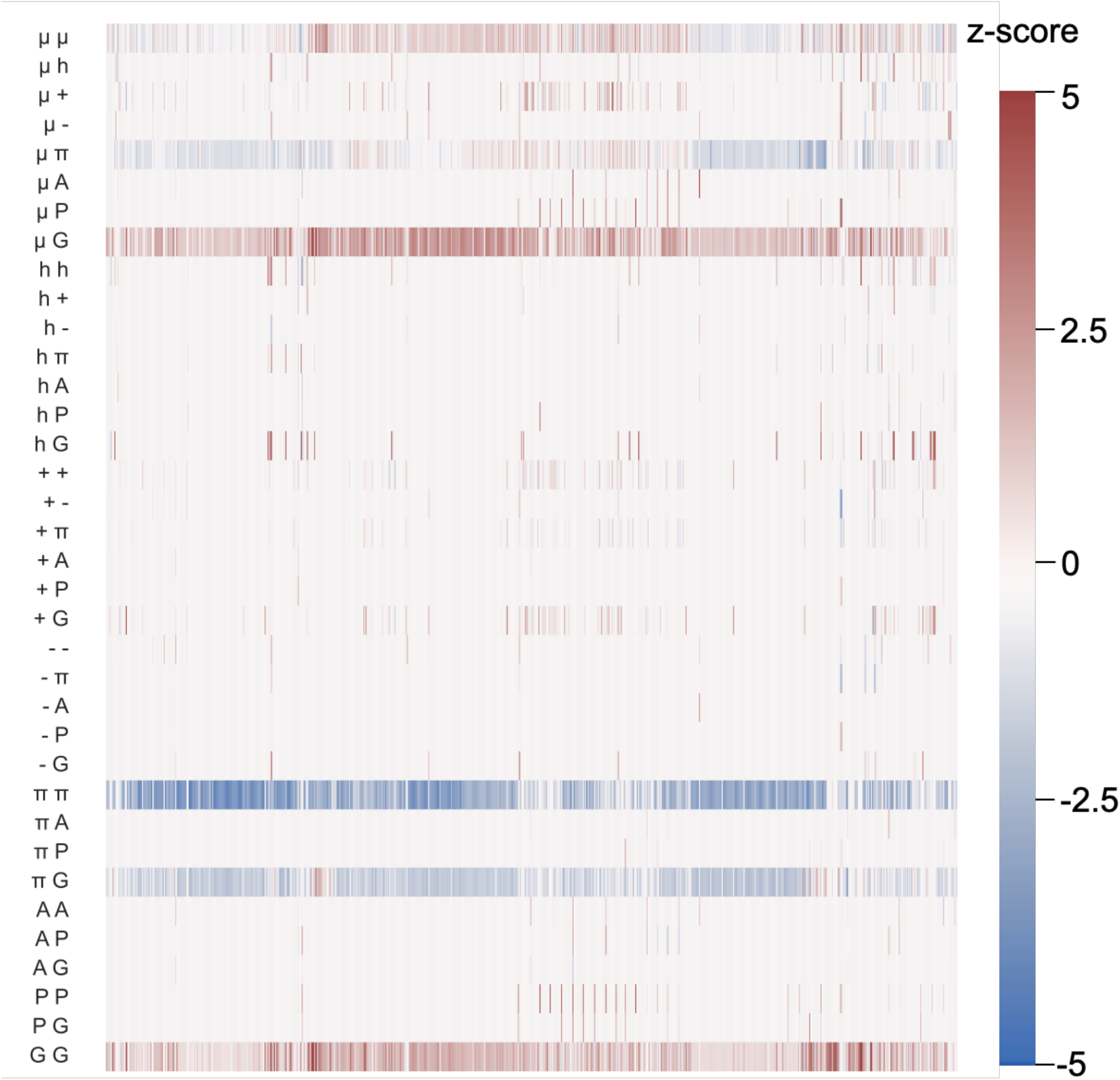
Elements of the z-score matrices for 849 homologs of A1-LCD. The color bar provides quantitative annotation for the heat map. This analysis reveals the following statistically significant binary patterns across homologs: (i) pronounced segregation of polar and Gly residues from one another (μG); (ii) uniform dispersion of aromatic residues with respect to one another (ππ); and (iii) segregation of Gly residues into clusters (GG).

### Analysis of C-terminal Domains (CTDs) in bacterial RNases E

In bacteria, the protein RNase E is a critical driver of the formation of the RNA degradasome ^41-43^. The architecture of this protein includes a conserved DEAD-box RNA helicase and a disordered C-terminal domain (CTD). In *C. crescentus*, the RNase E CTD is necessary and sufficient to drive phase separation. *In vivo*, RNase E drives the formation of cytoplasmic foci that colocalize with other exonucleases. This degradation body has been termed the Bacterial Ribonucleoprotein body or BR-body ^43^. The CTD of the *C. crescentus* RNase E has a blocky architecture characterized by linear segregation of oppositely charged residues ^42^. This leads to multivalence of oppositely charged blocks ^44^. Experiments suggest that this architecture is essential for the formation of BR bodies ^43^. In contrast, the *E. coli* RNase E, which lacks the blocky patterning of oppositely charged residues, does not form cytoplasmic condensates; instead, it forms membrane-tethered puncta *in vivo* and under the solution conditions that have been investigated to date, it does not form liquid-like condensates *in vitro* ^41,43^. Sequences of each CTD are shown in Supplementary Information (**Figure S2**).

We analyzed CTDs from *C. crescentus* and *E. coli* using NARDINI **(Figure 8)**. For the CTD of RNase E from *C. crescentus*, the patterns with the largest z-scores are in descending order: δ_h+_, δ_+–_, Ω_+_, δ_μ+_, δ_h–_, Ω_–_, Ω_h_, and δ_μ-_, respectively. In fact, the z-score values for all these patterns are greater than +5. This highlights statistically significant linear segregation of positively charged residues from all other residues, leading to a sequence with a multi-block architecture. Accordingly, the CTD of the *C. crescentus* RNase E may be classified as a blocky polyampholyte that also includes clusters of hydrophobic residues. In contrast, while the statistically significant segregation of positively charged residues is preserved in the CTD of RNase E from *E. coli*, the z-score values for δ_μ+_, δ_+-_, and Ω_+_ are at least two-fold smaller for this system when compared to those of the CTD of RNase E from *C. crescentus* (**Figure 8**). This quantitative difference in patterning preferences might explain the differences in driving forces for phase separation. Specifically, the overall weakening of the extents of linear segregation of oppositely charged residues as well as charged and hydrophobic residues from one another correlates with the observation of distinct phenotypes and driving forces for CTD mediated phase transitions of RNases E that control the formation of BR bodies.

**Figure 8.**
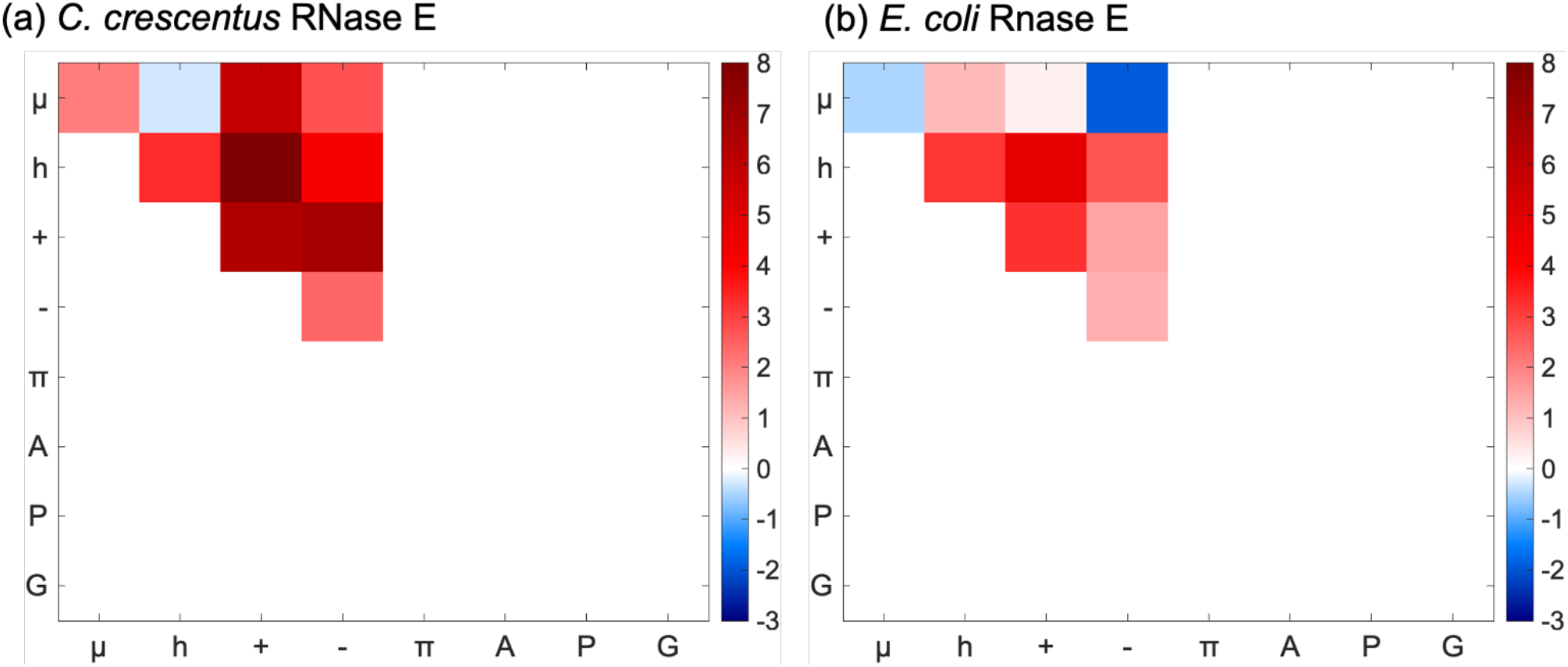
Direct comparison of z-score matrices of RNase E from (a) *C. crescentus* and (b) *E. coli*. Patterns associated with charged residues in *C. crescentus* RNase E (left) are > +2.4 standard deviations away from the null-scramble model in the positive direction. *E. coli* RNase E shows non-random segregation of positive residues and hydrophobic residues as well as from other residues, and hydrophobic residues also contribute to non-random patterns. Unlike the *C. crescentus* RNase E, patterns involving negative residues in *E. coli* RNase E do not significantly deviate from the null model.

Next, we analyzed z-scores for CTDs from 1084 orthologous RNases E (**Figure 9**). These sequences were excised from the RNases E of alpha and betaproteobacterial classes. In accord with the results summarized in **Figure 8**, there is a clear quantitative difference in statistically significant patterns between CTDs of RNases E drawn from the two classes. Sequences from the alphaproteobacterial class show a strong preference (high positive z-scores) for segregation of positively charged, hydrophobic, and polar residues into distinct clusters. This preference is weakened or eliminated for CTDs of RNases E drawn from the betaproteobacterial class. Taken together with published results for BR bodies, the results in **Figures 8** and **9**, point to how the quantitative modulation of binary patterns impact phase separation of bacterial RNases E.

**Figure 9:**
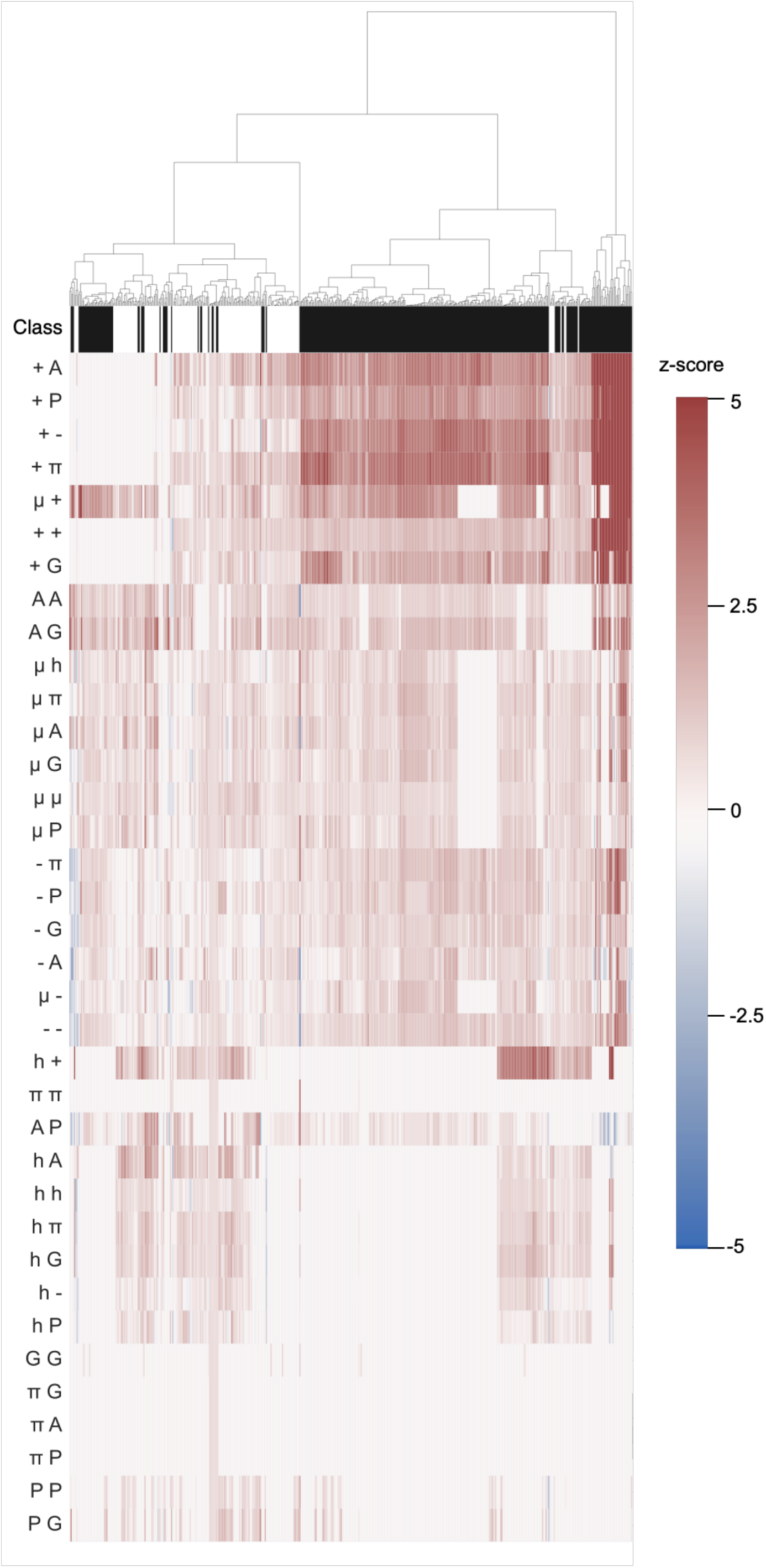
Analysis of z-score matrices across CTDs from 1084 RNase E orthologs. Each row below the row with labeled as Class denotes the z-score for a distinct binary pattern. The color bar provides annotation of the z-scores. Positive z-scores, denoted as red colors, quantify the extent of linear clustering of residue types within a sequence. Within the alphaproteobacterial class (black) there is a clear preference for the segregation of positively charged residues with respect to all other residue types leading to tracts of basic residues. This preference is weakened for CTDs of RNases E from the betaproteobacterial class (white). The class-specific preferences are illustrated using a dendrogram shown at the top of the figure. The dendrogram was generated using the Frobenius norms of z-score matrices, where the norms were used as Euclidean distances and Ward’s clustering was used to generate the dendrograms.

### Analysis of IDRs in bacterial single-stranded DNA binding proteins (SSBs)

Another example of an essential IDR in bacteria is the intrinsically disordered linker (IDL) in single-stranded DNA binding proteins (SSBs) that plays critical roles in bacterial DNA replication and repair. Modular architectures of SSBs include an ordered DNA-binding domain (OB-fold), followed by a hypervariable intrinsically disordered linker (IDL) that is connected to a conserved C-terminal tip ^45-47^. Recent work has also shown that the SSB from *E. coli* can mediate phase separation with DNA ^48^; however, the sequence requirements for these functionalities have yet to be elucidated. Using NARDINI, we sought to generate testable hypotheses regarding statistically significant binary patterns within SSB IDLs. We focused on the *E. coli* SSB IDL since it is the most well characterized both in terms of its impact on the cooperativity of ssDNA binding and phase behavior ^45-54^.

For the *E. coli* SSB IDL, the binary pattern that is statistically significant pertains to the linear segregation of Gly from all other residues. Gly residues are found in a series of short linear clusters, segregated from all other residues in the SSB IDL sequence. The blocks of Gly residues are often interspersed by Pro, giving rise to positive deviations of δ_PG_ from the null model (**Figure 10)**. These features are highlighted in the sequence of the IDL from *E. coli* SSB: TMQML**GG**RQS**GG**APA**GG**NI**GGG**QPQ**GG**W**G**QPQQPQ**GG**NQFS**GG**AQSRPQQSAPAAPSNEPP.

**Figure 10.**
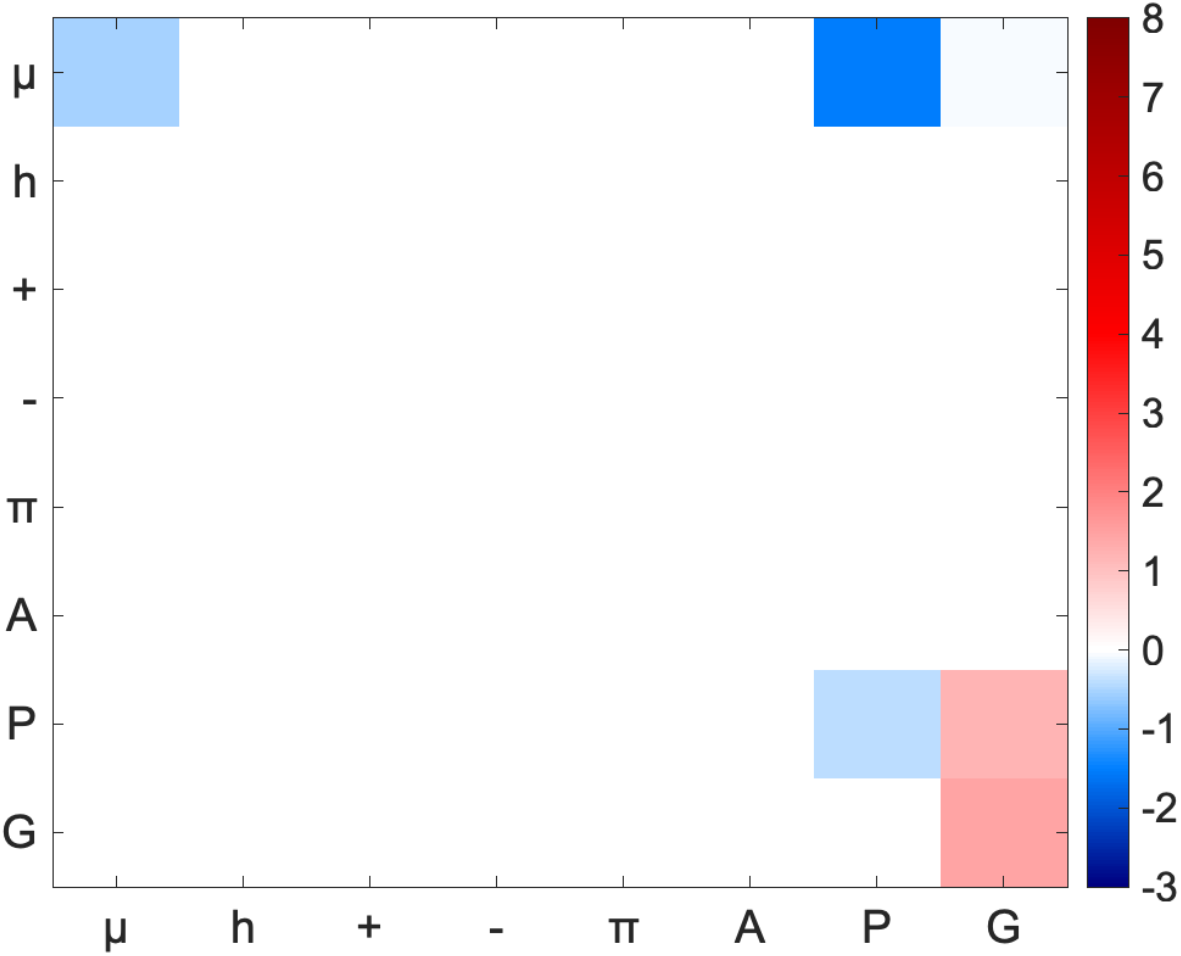
The z-score matrix of *E. coli* SSB IDL. The positioning of glycine, proline, and polar residues along the sequence is significant. The statistically significant deviations from the null model include the linear clustering of Gly residues and the punctuation of these clusters by Pro residues giving the IDL.

We assessed whether segregation of Gly residues is conserved across IDLs excised from 1523 SSBs of orthologous bacteria. The results are summarized in **Figure 11**. A large fraction of the IDLs feature clusters of Gly residues that are segregated from other residues. This is based on the z-scores we observe for Ω_G_. Additionally, these sequences also feature significant segregation of polar residues with respect to all residues with z(Ω_μ_) being greater than +1.5.

**Figure 11:**
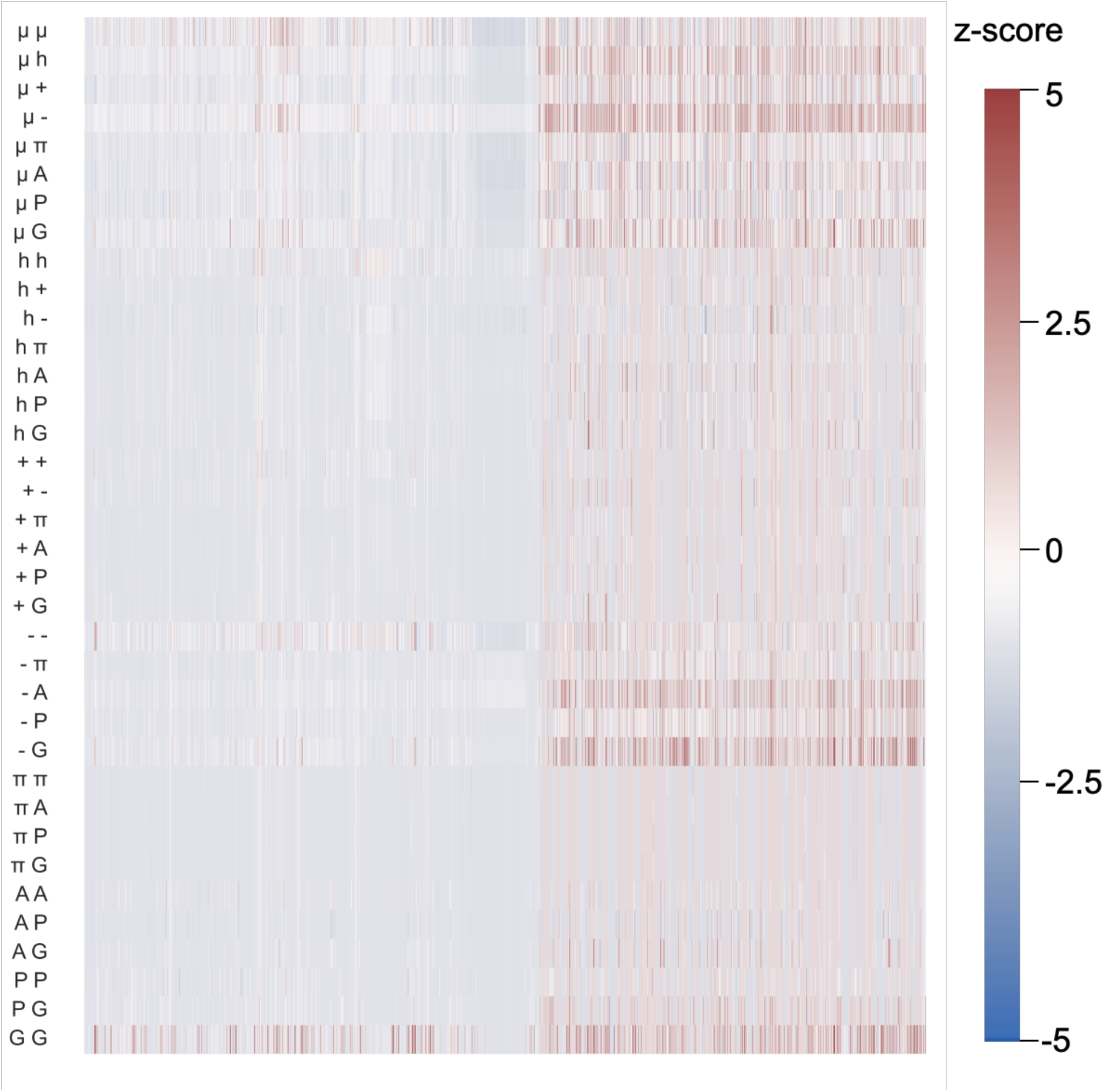
Heatmap of z-scores across IDLs of 1523 orthologous SSBs. Notice the positive z-scores for Ω_G_ labeled as GG in the figure. Other statistically significant patterns include the segregation of polar residues from Gly (μG) and acidic residues (μ-).

Next, we asked if the linear segregation of Gly and / or polar residues was a phylum- or class-specific feature. Classes that contained more than 5% of the total number of sequences were analyzed separately **(Figure 11)**. Statistically significant clustering of Gly residues was conserved across three out of the six evaluated classes: actinobacteria, α-proteobacteria, and γ-proteobacteria, whereas the IDLs of SSBs from bacilli, ε-protobacteria, and spirochaetia show minimal deviations from the null model for all binary patterns.

**Figure 11.**
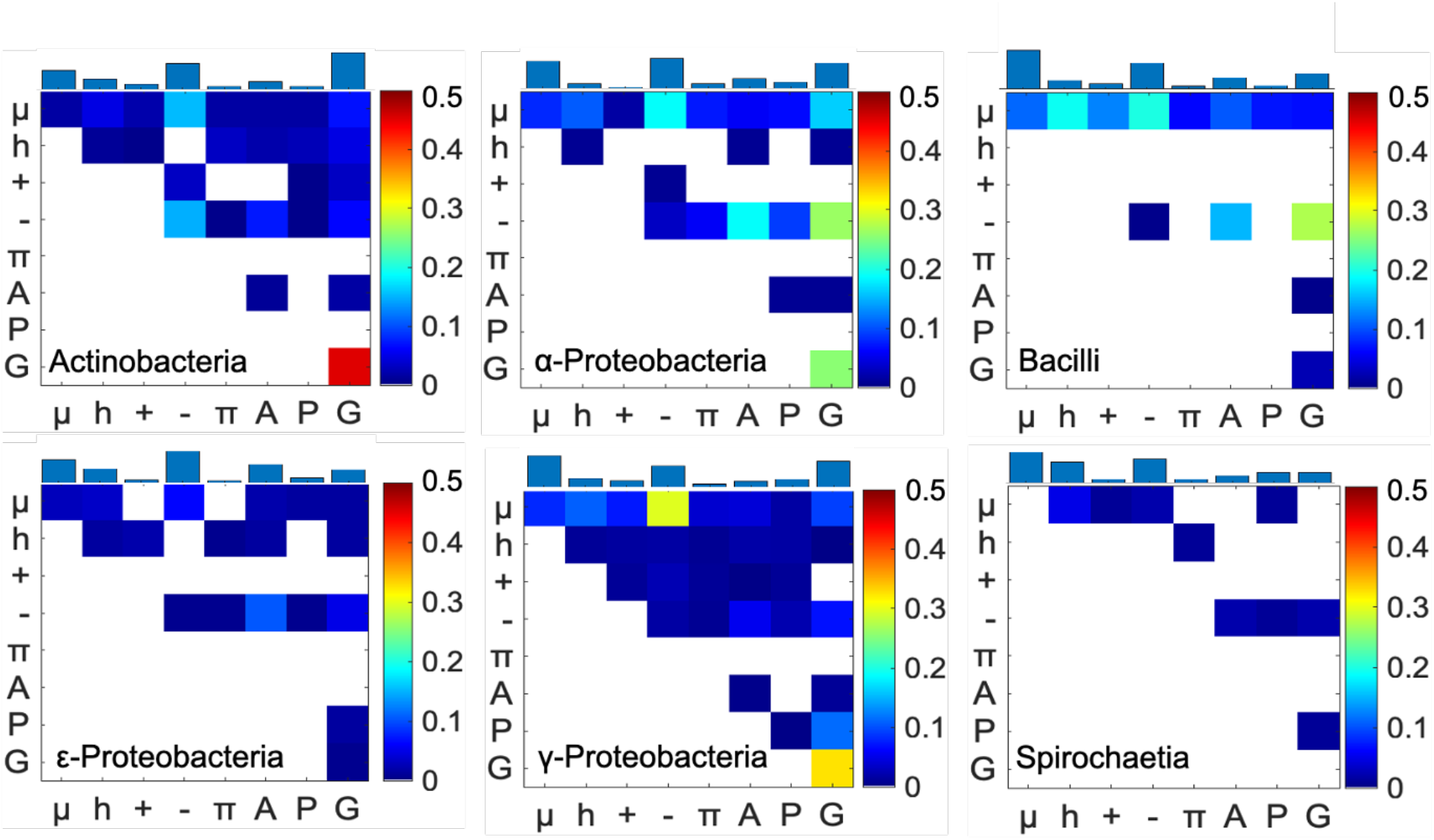
The frequency of observing a non-random pattern for the SSB IDLs. Squares within the checkerboard plot that do not rise about the 10% threshold are shown in white color. In all, we analyzed six phylum classes. These include actinobacteria (n = 190), bacilli (n = 239), γ- proteobacteria (n = 359), α-proteobacteria (n = 122), ε-proteobacteria (n =143), and spirochaetia (n = 101). The bar graph on the top of the matrix represents the relative frequencies of observing a non-random feature (z > +1.5) involving each residue / residue type.

The *E. coli* SSB IDL and IDLs from orthologous proteins have features that are reminiscent of elastomeric IDRs that are known to control elastic responses of materials such as extracellular matrices ^55-58^. Examples of such elastomeric IDRs include Gly-rich regions within resilin, as seen with repeats of PSSSYGAP**GGG**N**GG**R. Other examples include stretches such as P**G**Q**G**QQ from Q-rich proteins such as gluten ^59^, Gly, and Ser rich motifs in silk ^60^, motifs such as Y**G**H**GG**N/G in cell wall proteins of higher plants ^61^, and repetitive motifs such as F**GG**M**GGG**K**GG** from abductin, a protein that makes up hinge ligaments that control the swimming behaviors of mollusks ^62^. Further, if random binary patterns give rise to canonical Flory random coil behaviors, as they do for unfolded states of generic intrinsically foldable proteins ^63^, then the IDLs of SSBs would still be akin to elastic materials. This is because the free energies of such systems, which would be purely entropic in nature, is concordant with Hooke’s law^64^. Therefore, it appears that the IDLs of SSBs encode features that ensure elastomeric properties, either through specific patterning preferences or through minimal deviations from the null model. A direct test of this hypothesis would be to design sequences with different degrees of elastomeric responses, achievable using NARDINI, and test their impacts on ssDNA binding cooperativity as well as the phase behaviors of SSBs.

## Discussion

We have developed a method to uncover statistically significant binary patterns within a IDP sequences. Our work represents a generalization of approaches that have been brought to bear for identifying SLiMs and conserved features in IDRs ^13,65-67^. The method, referred to as NARDINI, is available as an extension for localCIDER ^28^, and can be deployed across orthologous systems.

Binary patterning parameters quantify the extent to which a residue, or a residue type, is positioned within the sequence with respect to other residues ^9,24,25,28,29^. We showed that the original versions of binary patterning parameters labeled κ ^29^ and Ω ^20,30^ and adaptations thereof cannot be compared across sequences of different compositions because the value of the patterning parameter that would be expected at random is inherently dependent upon the sequence composition ^28^. To solve this problem, we use the deviation (z-score) of the observed patterning parameter from a null model. Our method is a zeroth-order approximation of a uniform background that allows for comparisons of hypervariable sequences in a statistically meaningful way.

Our efforts were inspired by and build on the advances of Zarin, et al., who developed methods for discovering evolutionary signatures within IDRs ^4,5^. These signatures are defined as sequence features that show statistically significant deviations from a chosen null expectation. Zarin et al., ^4,5^ examined a total of 82 sequence features. This includes the κ_+-_ parameter as originally formulated by Das and Pappu ^29^. Other parameters of the overall feature vector include features of the fraction of charged residues, the isoelectric point, and the presence of motifs and repeats. We focused here on the development of a method that enables the extraction of statistically significant binary patterns within an IDP / IDR. Our method, as shown using three examples, can also be used to quantify the extent to which a specific binary pattern or patterns are conserved across homologs. Our focus on binary patterns was motivated by numerous reports that have documented the importance of these patterns in determining sequence-ensemble relationships and phase behaviors of IDPs / IDRs ^9,10,20,24-27,29,30,34,35,43,44,68-75^. An essential difference is that Zarin, et al., derive their null model expectations by explicitly simulating evolution of a sequence of interest based on the conservation of SLiMs with amino acid substitutions. This approach does not leave the amino acid composition fixed. While this approach is well suited for the goals pursued by Zarin et al., our approach, which generates a null model using a fixed composition, is intended for use in identifying statistically significant binary patterns within a given sequence and across sequences. This is necessary for guiding sequence designs and for interpreting measured and computed sequence-ensemble relationships. In ongoing work, we have found that the joint use of the evolutionary analysis of Zarin et al., and the NARDINI based identification of statistically significant binary patterns is instructive for large scale proteome wide analysis of IDRs. This joint approach leverages the distinctive features of the two methods viz., the evolutionary considerations and large numbers of features used by Zarin et al., and the composition specific z-score values for distinct patterns extracted using NARDINI.

In this work, we used the z-score method to identify binary patterns that show statistically significant deviations from null models for three different systems. The significant patterns we identify for the A1-LCD are consistent with findings from experiments ^20^. Similarly, the statistically significant linear segregation of positively charged residues from all other residue types and the blocky architecture for CTDs of bacterial RNases E is consistent *in vivo* phenotypes showing the influence of disrupting this architecture on the nature of RNA degradasomes. In *C. crescentus*, this body is a biomolecular condensate that exhibits liquid-like properties, whereas, in *E. coli*, it is a membrane-bound punctum ^43^. This discrepancy has been shown to be dictated by differences in the CTD sequence. In the CTD of *C. crescentus* RNase E, the blocky charge architecture of this sequence is a feature that significantly deviates from the null expectation. While the positively charged residues of the *E*. coli CTD are non-randomly positioned (Ω_+_), the negative residues are randomly positioned in relation to positively charged residues (δ_+–_) as well as in relation to other groups of residues (Ω_–_).The non-random segregation of charged residues (δ_+–_) is a feature that is consistent with over 27% of RNase E CTDs, and this could be a feature that is relevant for RNA binding within the degradasome.

In the IDL of *E. coli* SSB, we observed a conserved segregation of Gly residues (Ω_G_) whereby they are frequently found in short linear clusters along the sequence. We hypothesize that this feature could have implications for the cooperative binding of SSBs to single-stranded DNA and for the driving forces for phase-separation of SSBs ^45,46,48-50,52-54^. SSBs form homo-tetramers that generate a tetra-valent system to coordinate interactions with SSB interacting proteins ^47,52^. The *E. coli* SSB tetramer binds cooperatively to single-stranded DNA in different binding modes where the binding modes are classified by the number of nucleotides that are occluded by individual tetramers ^50,52,53^. Cooperativity of single-stranded DNA binding is governed by sequence features of the IDL ^45^. Specifically, cooperativity is enhanced when the IDL has features that are akin to low complexity domains enriched in polar amino acids, primarily Pro, Gln, and Gly ^76-79^. Conversely, cooperativity is diminished for long IDLs enriched in charged residues ^45,46^. The enrichment of Pro, Gln, and Gly that appear to govern cooperativity and their patterning were observed to be significant non-random patterns in the z-score analysis. Further, a recent study has shown that in response to DNA damage, membrane associated SSBs form condensates at the sites of DNA damage ^48^. These condensates are multicomponent bodies and concentrate other factors that contribute to DNA processing and metabolism. *In vitro* studies showed that the IDL is essential for driving the formation of liquid-like condensates. It could be that the patterning of Gly residues plays an important role in condensate formation – a feature that would be true of elastomeric sequences ^58,80^.

NARDINI is intended for use in *de novo* design of IDPs / IDRs with a view toward uncovering sequence-ensemble-function relationships and for identifying features that are likely selected for among functional orthologs. Our findings point to the possibility of using pattern-specific z-scores as order parameters for describing sequence-ensemble relationships and phase behaviors of IDPs / IDRs ^81,82^. Finally, although our analysis focused on binary patterns, there may be higher-order correlations that are determinants of sequence-function relationships encoded by hypervariable IDRs. The topic of higher-order correlations in sequence patterns is a focus for continued development of NARDINI.

## Acknowledgments

We thank Kiersten Ruff for helpful comments and critical analysis of the codebase as well as results generated using NARDINI. We are grateful to Kiersten Ruff and Mina Farag for critical reading of the manuscript. The source code is available as part of version 3.0 of localCIDER that is distributed through the Pappu lab Github. Version 3.0 of CIDER was written by Alex Keeley, a summer 2020 programmer in the Pappu lab. This work was supported by grants from the US National Science Foundation (MCB 1614766) and the US National Institutes of Health (5R01NS056114). Min Kyung Shinn is supported in part by the Center for Science & Engineering of Living Systems (CSELS) in the McKelvey School of Engineering at Washington University in St. Louis as a CSELS postdoctoral fellow.

