## Supplemental Material for "Uncovering non-random binary patterns within sequences of intrinsically disordered proteins"

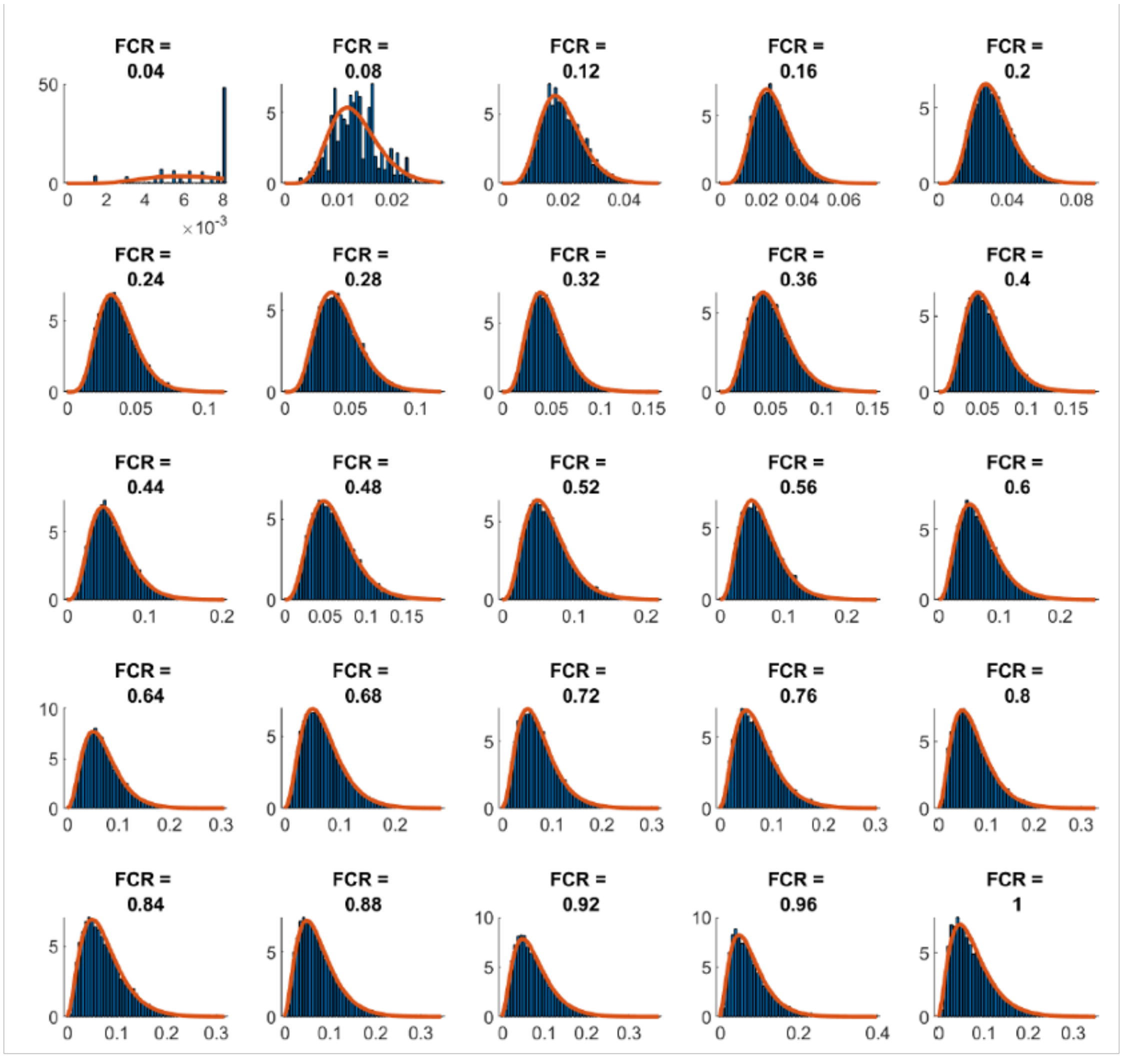


**Figure S1. Goodness of fit of the gamma distribution as a function of the fraction of the residues of interest for $\kappa_{+-}$.** Three different amino acids, A, E, and K were used for sequences of 50 residues. The residues of interest for $\kappa_{+-}$ are the charged residues, E and K. The net charge per residue for each sequence was fixed at 0, while the fraction of charged residues (FCR) was varied.

**hnRPA1**
FCR = 0.122 | NCPR = 0.049

MSYYHHHHHH LESTSLYKKA GFENLYFQGS MASASSSQRG RSGSGNFGGG
RGGGFGGNDN FGRGGNFSGR GGFGGSRGGG GYGGSGDGYN GFGNDGSNFG
GGGSYNDFGN YNNQSSNFGP MKGGNFGGRS GGSGGGGQYF AKPRNQGGYG
GSSSSSSYGS GRRF

<u>*E. coli* RNase E</u>
FCR = 0.346 | NCPR = 0.025

RQDGVRCVIV PNDQMETPHY HVLRVRKGEE TPTLSYMLPK LHEEAMALPS
EEEFAERKRP EQPALATFAM PDVPPAPTPA EPAAPVVAPA PKAAPATPAA
PAQPGLLSRF FGALKALFSG GEETKPTEQP APKAEAKPER QQDRRKPRQN
NRRDRNERRD TRSERTEGSD NREENRRNRR QAQQQTAETR ESRQQAEVTE
KARTADEQQA PRRERSRRRN DDKRQAQQEA KALNVEEQSV QETEQEERVR
PVQPRRKQRQ LNQKVRYEQS VAEEAVVAPV VEETVAAEPI VQEAPAPRTE
LVKVPLPVVA QTAPEQQEEN NADNRDNGGM PRRSRRSPRH LRVSGQRRRR
YRDERYPTQS PMPLTVA

<u>*C. crescentus* RNase E</u>
FCR = 0.406 | NCPR = -0.085

TGVLEGTTHV CEHCEGTGRV RSVESSALAA LRAVEAEALK GSGSVILKVS
RSVGLYILNE KRDYLQRLLT THGLFVSVVV DDSLHAGDQE IERTELGERI
AVAPPPFVEE DDDFDPNAYD DEEEEDDVIL DDEDDTDRED TDDDDATTRK
SARDDERGDR KGRRGRRDRN RGRGRRDERD GETESEDEDV VAEGADEDRG
EFGDDDEGGR RRRRRGRRGG RRGGREDGDR PTDAFVWIRP RVPFGENVFT
WHDPAALVGG GESRRQAPEP RVDAATEAAP RPERAEREER PGRERGRRGR
DRGRRQRDEA PVAEMTSVES ATVEAAEPFE APILAPPVIA GPPADVWVEL
PEVEEAPKKP KRSRARGKKA TETSVEAIDT VTEVAAEAPA PETAEPEAVE
VAPPAPTVEA APEPGPVVEA VEEAQPAEPD PNEITAPPEK PRRGWWRR

**Figure S2. Sequences of the hnRPA1 and RNases E CTDs from *E. coli* and *C. crescentus*.** Residues are color-coded as following: negatively charged (red), positively charged (blue), polar (green), hydrophobic (black), prolines (magenta), and aromatic (orange). The blocky architecture of *C. crescentus* RNase E is shown by the enrichment of blue or red stretches within the sequence. The fraction of charged residues (FCR) and the net charge per residue (NCPR) are noted. Both sequences are highly charged and polyampholytic where the net charge per residue approaches 0.
